# Comparative Evaluation of Adeno-Associated Virus and Lentivirus Mediated Gene Transfer in Adult Rat Optic Nerve

**DOI:** 10.64898/2026.05.12.724624

**Authors:** Chaimaa Kinane, Rajeshwari Koilkonda, Javier Gomez, Moxa Panchal, Tamdan Khuu, Venu Talla, Kevin K. Park

## Abstract

**Background:** The optic nerve serves as a vital conduit for visual signaling, and its degeneration in optic neuropathy results in irreversible vision loss. It is also a widely used model for studying central nervous system (CNS) injury and repair. Although adeno-associated virus (AAV) and lentivirus are extensively applied in CNS research, their transduction efficiency and cell-type specificity within the optic nerve remain poorly characterized. This study aimed to identify the most effective viral vector, serotype, and promoter for direct gene delivery to the adult rat optic nerve.

**Methods:** Sprague–Dawley rats (7–10 weeks) received intra-optic nerve injections of lentiviral or AAV vectors encoding GFP under different promoters (CAG, CMV, or GFAP). Two to three weeks post-injection, optic nerves were collected for immunohistochemistry with markers of oligodendrocytes (Olig2), astrocytes (GFAP, Sox9), and microglia (IBA1). Transduction efficiency and cell-type specificity were assessed using confocal microscopy.

**Results:** AAV2, AAV5, and lentivirus showed minimal transduction, with only sparse GFP-positive cells observed near injection sites. In contrast, AAV-PHP.eB carrying the CAG promoter yielded robust and widespread GFP expression near the injection site. Quantitative analysis revealed that approximately 90% of transduced cells were Olig2-positive oligodendrocytes, indicating strong tropism for this glial population.

**Conclusion:** AAV-PHP.eB driven by the CAG promoter enables efficient gene delivery to the optic nerve, with a predominant tropism for oligodendrocytes. This targeted intra-optic nerve injection approach offers a reliable platform for manipulating oligodendrocytes and investigating mechanisms of CNS development, injury, and repair relevant to both optic neuropathies and other CNS diseases.

## INTRODUCTION

The optic nerve plays a fundamental role in vision by transmitting visual information from the retina to the brain. Damage to the optic nerve, as seen in diseases such as glaucoma, optic neuritis and certain forms of retinal degeneration, can result in irreversible vision loss and ultimately blindness.^1–6^ Understanding the cellular and molecular underpinnings of these conditions is essential for developing effective therapies. One of the most powerful approaches for investigating gene function and disease mechanisms in the central nervous system (CNS), including the optic nerve, is viral-mediated gene delivery. Viral vectors, particularly adeno-associated viruses (AAVs) and lentiviruses, have become indispensable tools in neuroscience due to their ability to deliver genetic material with relatively high specificity and safety profiles.^7–11^

To date, extensive studies have examined the tropism and transduction efficiency of various AAV serotypes in the brain and spinal cord.^12–17^ These investigations have identified serotype specific patterns of cell targeting, with certain AAVs showing strong preferences for neurons, astrocytes, or oligodendrocytes depending on the route of administration and choice of promoter. ^7–9,18^ Similarly, lentiviruses have been explored for their ability to stably integrate into the host genome and target a broad range of dividing and non-dividing. ^19–21^ However, despite this growing body of research in other parts of the CNS, there remains a notable gap in our understanding of viral vector performance in the optic nerve.

Compared to the brain and spinal cord, the optic nerve shares certain structural and cellular similarities but also exhibits key differences that are critical to consider when evaluating gene delivery strategies. Like the spinal cord, the optic nerve contains glial cell populations such as myelinating oligodendrocytes, astrocytes, and microglia, as well as a dense extracellular matrix. ^22–24^ These common features suggest that insights from viral vector studies in the spinal cord may offer some guidance for approaches in the optic nerve. However, a difference lies in the absence of neuronal cell bodies within the optic nerve. Unlike the spinal cord, which contains both neurons and glia, the optic nerve consists primarily of axons originating from retinal ganglion cells (RGCs), whose cell bodies reside in the retina. These anatomical and cellular differences emphasize the need for direct, systematic evaluation of viral vectors in the optic nerve, rather than relying on data derived from other CNS tissues. Identifying optimal AAVs for gene delivery to the optic nerve is crucial not only for advancing therapies for optic neuropathies but also for expanding the utility of optic nerve injury models, which are widely used to study CNS injury responses.^25,26^ The optic nerve provides a practical and accessible surgical target in animal models, offering a consistent and reproducible platform compared to more complex brain or spinal cord procedures.^26–28^ In this study, we assessed the efficiency and cellular tropism of two major viral platforms—lentivirus and AAV—following direct injection into the adult rat optic nerve. By comparing different viral constructs, serotypes, and promoter configurations, we sought to identify the conditions that yield robust gene expression in the optic nerve.

## METHODS

### Ethical Review

All animal procedures were conducted in compliance with protocols approved by the Institutional Animal Care and Use Committee (IACUC) at University of Texas Southwestern Medical Center (UTSW) and in accordance with the Association for Research in Vision and Ophthalmology Statement for the Use of Animals in Ophthalmic and Vision Research.

### Animals

Adult male Sprague-Dawley rats (*Taconic Bioscience Model No. SD-M, NTac:SD*) were used in this study. The rats were 7 to 10 weeks old. Rats were housed in individually ventilated cages under a 12:12 hour light/dark cycle. Food and water were supplied *ad libitum*. All cages included nesting material and shelters to provide environmental enrichment.

### AAVs and Lentivirus

AAVs and lentiviruses except for AAV2-Cytomegalovirus (CMV)-GFP were purchased from SignaGen Laboratories. These viruses were supplied at the following titers: AAV-PHP.eB-glial fibrillary acidic protein (GFAP)-GFP (Catalog # SL116362; Titer: 2.19 ×10^13^ viral genome (vg)/mL); AAV-PHP.eB-early enhancer/chicken β-actin (CAG)-GFP (Catalog # SL116010, 1.69 ×10^13^ - 2.58 ×10^13^ vg/mL); AAV5-CAG-GFP (Catalog # SL100823, 2.08 ×10^13^ vg/mL; AAV5-GFAP-GFP (Catalog # SL101519; 3.52 ×10^13^ vg/mL); LV-CAG-GFP (Catalog # SL100270; 1.46 ×10^9^ transducing unit (TU)U/ml). AAV2-CMV-GFP (0.9 × 10^13^ vg/ml) was produced at the UTSW Ophthalmology Gene Editing and Viral Production Core. Lentivirus titers are reported as TU/ml, whereas AAV titers are expressed as vg/ml, reflecting functional infectivity and physical particle number, respectively.

### Intra-Optic Nerve Injection

Although intra-optic nerve injection has been attempted, achieving precise, controlled, and minimally invasive delivery remains technically challenging due to the small size, position and fragility of the optic nerve. ^29,30^ To address these limitations, we developed a refined injection approach using a fine glass micropipette coupled with a nanoliter injector, enabling controlled and localized delivery of viral vectors directly into the optic nerve. (Supplementary Fig. S1).

Adult rats were anesthetized with an intraperitoneal injection of Ketamine/Xylazine (50 mg/ml and 5 mg/ml). The conjunctiva was incised and gently contracted, and the overlaying orbital fat and connective tissue were carefully separated using forceps to expose the optic nerve. Extreme care was taken to avoid any damage to the ophthalmic artery. Prior to injection, viral aliquots were thawed on ice, and Fast Green FCF (Cat. #25053-02; Electron Microscopy Sciences) was added at a 10% concentration to aid visualization of the virus during the procedure. Once the optic nerve was exposed, pulled glass micropipettes with a small tip diameter (< 1 μm) connected to a nanoinjector (Nanoliter 2020 injector head, World Precision Instruments) were used to deliver the viral vectors into the optic nerve. The pipette was inserted into the exposed optic nerve under the stereomicroscope (Prescott’s Microscope, Fixed Post Table Top system, USA, approximately 2– 3 mm posterior to the optic disc (Supplementary Fig. S1). Approximately 300 nl of each virus was injected at a rate of 2 nl/second to minimize pressure-induced damage. Thus, for lentivirus, about 4.38×10^5^ TU was injected per optic nerve whereas for AAV, following amount of vectors were delivered for each optic nerve; AAV-PHP.eB-GFAP-GFP at 6.57 × 10^9^ vg, AAV-PHP.eB-CAG-GFP at 5.07 × 10^9^ vg, AAV5-CAG-GFP at 6.24 × 10^9^ vg, AAV5-GFAP-GFP: 1.06 × 10^10^ vg; AAV2-CMV-GFP at 2.07 × 10^9^ vg was injected per optic nerve. After the procedure, animals received a subcutaneous injection of sustained release-Buprenorphine for analgesia and were monitored for signs of ocular infection, hemorrhage, or distress.

### Tissue Processing

Animals were humanely euthanized in accordance with the American Veterinary Medical Association guidelines by CO_2_ inhalation followed by a confirmatory secondary method. Two to three weeks post-injection of lentivirus or AAVs, animals were then transcardially perfused with ice-cold phosphate-buffered saline (PBS, pH 7.4), followed by 4% paraformaldehyde (PFA) in phosphate-buffered saline (PBS, 1X). Eyes with optic nerves attached were carefully dissected and post-fixed for 2-3 hours in 4% PFA at room temperature (RT). Tissues were washed three times with 1× PBS (1X) and transferred to 30% sucrose in PBS for cryoprotection, remaining at 4°C overnight until they sank.

Optic nerves are then embedded in optimal cutting temperature (OCT), frozen on dry ice, and stored at −80 °C until use. Optic nerves were sectioned longitudinally at 10 μm using a cryostat (Leica CM3035) and mounted on Superfrost Plus slides. Slides were air-dried for about 30 minutes and stored at –20 °C until immunostaining.

### Immunohistochemistry

Sections were permeabilized in PBS containing 0.3% Triton X-100 (Cat. #T8787, Millipore Sigma), and nonspecific binding was blocked with 3% bovine serum albumin (BSA) in PBST for 1 hour at RT. Primary antibodies were diluted in blocking solution, combined as indicated, and applied overnight at 4 °C: Chicken anti-GFAP (1:1000, ab4674, Abcam), Rabbit anti-Olig2 (AB9610, Millipore), Goat anti-Sox9 (1:200, AF3075-SP, R&D), and Goat anti-IBA1 (1:500, 019-19741, FUJIFILM Wako). Slides were washed three times in PBST, then incubated for 1 hour at RT with species-appropriate secondary antibodies: Alexa Fluor 594 goat anti-chicken (1:500, Jackson ImmunoResearch), Alexa Fluor 647 goat anti-rabbit (1:500, Abcam), and Alexa Fluor 647 donkey anti-goat (1:500, ThermoFisher). After incubation, slides were mounted with Fluoromount-G (Cat. #00-4959-52, Invitrogen), coverslipped, and imaged using a Zeiss LSM 780 confocal or a Keyence BZX-800 microscope.

### Quantification and Statistics

Cell transduction was determined by systematic cell by cell colocalization analysis using confocal microscopy (Zeiss 780 Inverted). For each animal, optic nerve sections encompassing the injection site were imaged at high resolution (z stacks, 0.5 µm intervals). GFP^+^ cells within each field were identified by fluorescence colocalization with DAPI^+^ nuclei and manually counted using Image J (Fiji) with the cell counter plugin. Co-expression of Olig2, Sox9, or IBA1 was determined by channel overlap. Quantification was performed within a defined region of interest (ROI) of consistent size applied across all images. For each animal, 3-4 sections were analyzed, and values were averaged per animal prior to comparison across groups. Transduction specificity was calculated as the percentage of GFP^+^ cells co-expressing each marker relative to the total GFP^+^ population, with each GFP^+^ cell individually assessed for colocalization with lineage-specific markers.

Statistical analyses were performed using GraphPad Prism 10, with data presented as mean ± SEM. One-way ANOVA followed by Tukey’s post hoc test was used, and significance was defined as *p* < 0.05 (* *P* < 0.05, ** *P* < 0.01, *** *P* < 0.001, and **** *P* < 0.0001).

## RESULTS

### Comparative Analysis of AAV2, AAV.PHP.eB, and AAV5 Transduction Efficiency Driven by Ubiquitous (CAG) or GFAP Promoters

We evaluated the gene delivery efficiency of several commonly used AAV serotypes in the adult rat optic nerve. Specifically, we assessed AAV2 carrying a CMV promoter and AAV5 vectors containing either a ubiquitous CAG promoter or a GFAP promoter. We also assessed AAV-PHP.eB with a GFAP promoter. The viral vectors and promoter combinations used in this study were intentionally chosen to represent AAV serotypes and promoters with distinct, well-documented transduction properties in the central nervous system (CNS), particularly in the context of optic nerve and glial versus neuronal targeting.

AAV2 was included because it is one of the most widely used and extensively characterized serotypes in the CNS and visual system, with a long history of application in retinal gene delivery studies. Its use provides an important benchmark for comparison with newer or alternative capsids. AAV5 was selected based on previous reports demonstrating its ability to efficiently transduce astrocytes and glial populations in the optic nerve when injected intravitreally near the optic disc paired with GFAP promoters. AAV-PHP.eB, a recently engineered capsid, was included due to its strong CNS tropism and its demonstrated ability to robustly transduce CNS cells following either local or intravenous administration, making it of particular interest for optic nerve targeting.

The promoters chosen (CMV, CAG, and GFAP) similarly reflect commonly used elements with well-established profiles. CMV and CAG are widely used pan-cellular promoters that enable strong and broad transgene expression, whereas the GFAP promoter was selected to assess astrocyte-enriched transduction patterns. Each vector encoded GFP and was injected directly into the optic nerve approximately 2–3 mm distal to the optic disc. Optic nerves were collected about three weeks post-injection for immunohistochemical analysis. We found that these three AAVs assessed (i.e., AAV2-CMV-GFP, AAV-PHP.eB-GFAP-GFP, AAV5-CAG-GFP, and AAV5-GFAP-GFP) resulted in minimal transduction in the optic nerve tissue (Fig. 1A). Notably, most cells that appeared to exhibit GFP signal were in fact autofluorescent, indicating the absence of true GFP expression (Fig. 1B and Supplementary Figure 2). These cells are likely macrophages accumulated at the injection site, which are known to display autofluorescence under conventional epifluorescence microscopy. Only rare GFP-positive puncta lacking signs of autofluorescence (i.e., no red Cy3 signal), indicative of true transduced cells, were observed near the injection site (Fig. 1A). This pattern was consistent across all biological replicates. Notably, even with the broadly active CMV or CAG promoter, these AAVs did not achieve meaningful transduction in optic nerve glial cells. These results demonstrate that both AAV2 and AAV5 vectors, when delivered locally, have limited ability to deliver genes to cells in the adult optic nerve.

**Figure 1.**
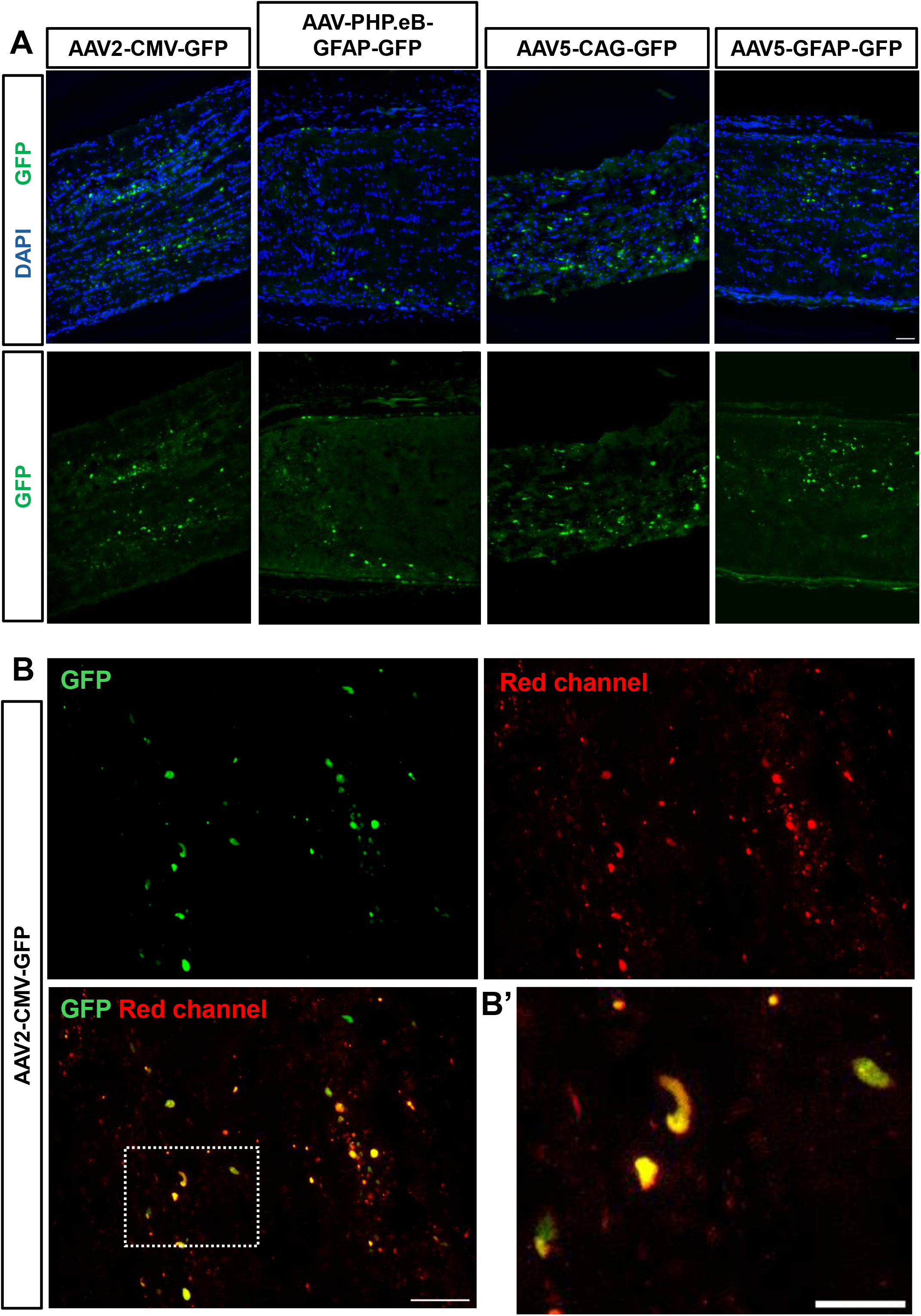
Assessment of AAV serotypes and promoters for transgene delivery to optic nerve cells following intra-optic nerve injection. (A) Representative longitudinal optic nerve sections following optic nerve injection of AAV2-CMV-GFP, AAV-PHP.eB-GFAP-GFP, AAV5-CAG-GFP, and AAV5-GFAP-GFP. (B) A higher magnification image of an optic nerve with AAV2-CMV-GFP. GFP signal is shown in the green channel. A red channel without any immunostaining was included to identify autofluorescent cells, which exhibit overlapping signals in the red and green channels. AAV2-CMV-GFP, N= 2; AAV-PHP-GFAP-GFP, N= 4; AAV5-CAG-GFP, N=4; AAV5-GFAP-GFP, N= 2. Scale bars, 50 μm.

### Lentivirus mediates transduction in a subset of cells, although overall efficiency is limited

We next evaluated the performance of lentivirus using the same local injection paradigm. Lentiviral vectors encoding GFP under the control of the CAG promoter were injected into the optic nerve and analyzed two weeks later. A small number of GFP-positive cells were detected within the optic nerve (Fig. 2). Transduced cells were immunoreactive for Sox9 (an astrocyte marker; Fig. 2C) or IBA1 (a microglial marker; Fig. 2D), indicating that lentiviral vectors driven by the CAG promoter can mediate gene delivery to these two cell types in the optic nerve, albeit with low efficiency. Reporter expression was also observed in the optic nerve sheath near the injection site (Fig. 2A). Thus, although lentiviral transduction efficiency exceeded that of the initial AAVs tested above, it remained substantially low. Together, these results indicate that lentivirus provides some but only limited transduction efficiency in the adult optic nerve following direct parenchymal injection.

**Figure 2.**
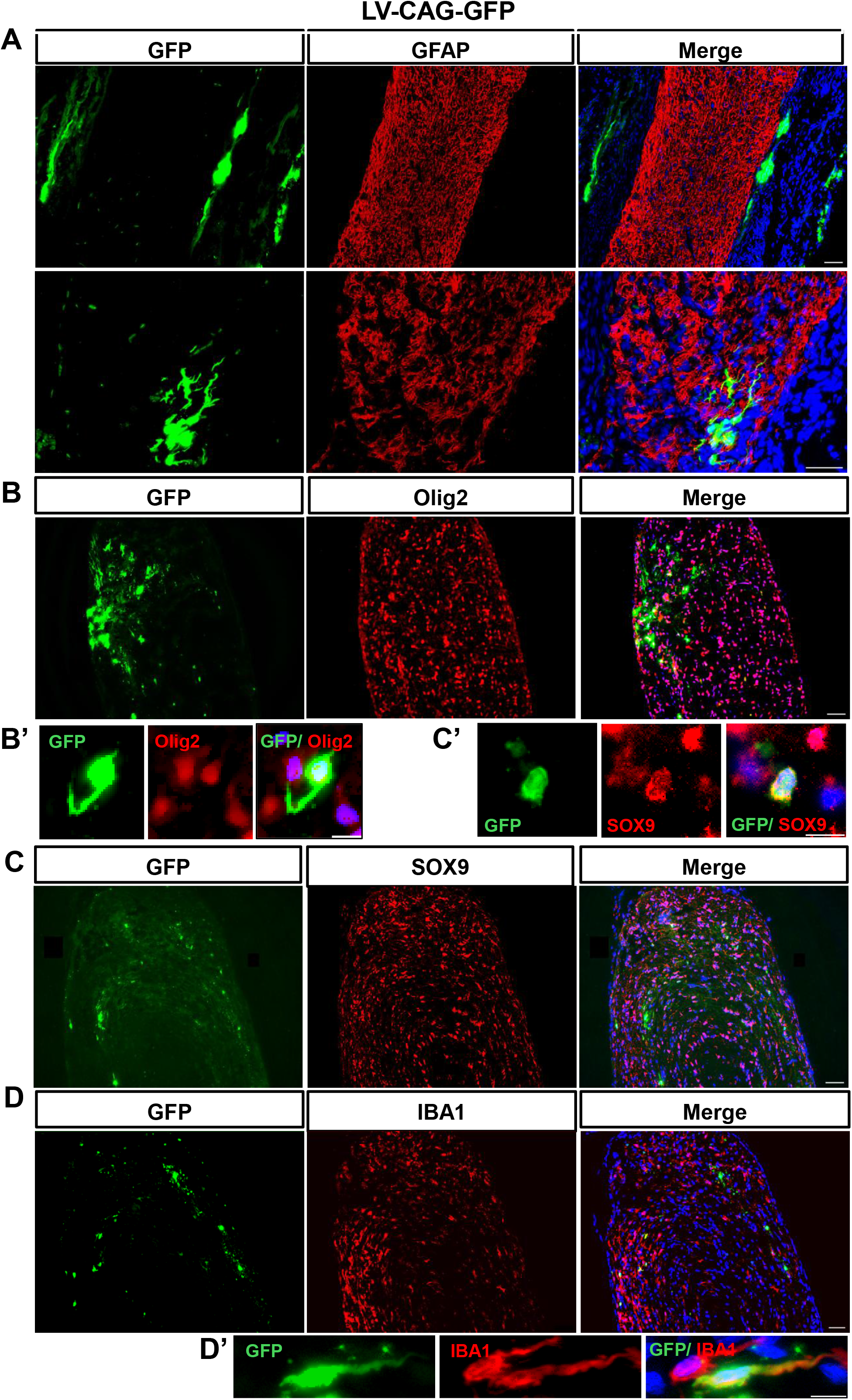
Intra-optic nerve injection of lenti-CAG-GFP results in gene delivery in a few cell types. Representative images showing GFP expression (green) following lenti-CAG-GFP injection into the optic nerve. (A) Immunostaining with a GFAP antibody showing two representative optic nerve sections. The upper panel displays images acquired at 20× magnification, while the lower panel presents higher-magnification views. (B) Immunostaining with an Olig2 antibody. (B’) A higher magnification image showing GFP and Oligo2-immunoreactive cells. (C) Immunostaining with a Sox9 antibody. (C’) A higher magnification cropped image showing GFP and Sox9-immunoreactive cells. (D) Immunostaining with an IBA1 antibody. (D’) A higher magnification of showing GFP and IBA1-immunoreactive cells. N= 4. Scale bars, 20 μm (A, upper panel); 50 μm (A, lower panel), and 10 μm in higher-magnification cropped images (B’, C’, and D’).

### AAV-PHP.eB with a CAG Promoter Achieves Strong Transgene Expression

AAV PHP.eB is a rationally engineered variant of the AAV9 capsid that was generated through in vivo directed evolution. Specifically, a short peptide insertion was introduced into an exposed variable region of the AAV9 capsid (variable region VIII), followed by iterative selection for enhanced transduction efficiency within the central nervous system. This modification significantly increases capsid interaction with host endothelial and neural cell entry pathways, resulting in substantially improved CNS tropism compared with unmodified AAV9 in rodents. In contrast to the poor transduction seen with AAV2, AAV5, and lentivirus, injection of AAV-PHP.eB encoding GFP under the CAG promoter resulted in strong gene expression in the optic nerve. Non-autofluorescent GFP signal was clearly visible along the longitudinal axis of the nerve in most sections. GFP expression was observed across multiple consecutive optic nerve sections, indicating successful local spread of viral transduction beyond the immediate injection site. However, the number and density of GFP-positive cells progressively tapered toward the distal edges of the optic nerve (Figs. 3 and 4). GFP-positive cell coverage was concentrated near the injection site, with diminishing signal intensity and fewer GFP-positive cells at greater distances. Qualitatively, immunohistochemical assessment shows that most GFP-positive cells co-expressed Olig2 (Fig. 3A, 3B), indicating that oligodendrocytes are the primary target population for AAV-PHP.eB in the optic nerve under these conditions. GFP-positive cells were only rarely found to be immunoreactive for Sox9 or IBA1 (Fig. 4A–C). Quantification of transduced cell types shows that over 90% of GFP^+^ cells are positive for Olig2, whereas only a small fraction co-expressed Sox9 (~6%) or IBA (~1-2%) (Fig. 5A) indicating preferential targeting of oligodendrocyte-lineage cells. This pattern was further supported by cell density measurements, which revealed minimal GFP^+^/Sox9^+^ and GFP^+^/IBA1^+^ double positive cells, and a strong co-expression between GFP^+^ and GFP/Olig2 populations (Fig. 5B-D). Quantification of the GFP-positive areas revealed that within a cryosection with the highest GFP^+^ cell density, the coverage spanned approximately 0.1 mm^2^. These findings indicate that while viral transduction extends across several optic nerve sections, the effect remains largely localized rather than diffusely distributed throughout the entire nerve. Taken together, these results identify AAV-PHP.eB as an effective vector for gene delivery to glial cells in the adult optic nerve, with particularly strong tropism for oligodendrocytes.

**Figure 3.**
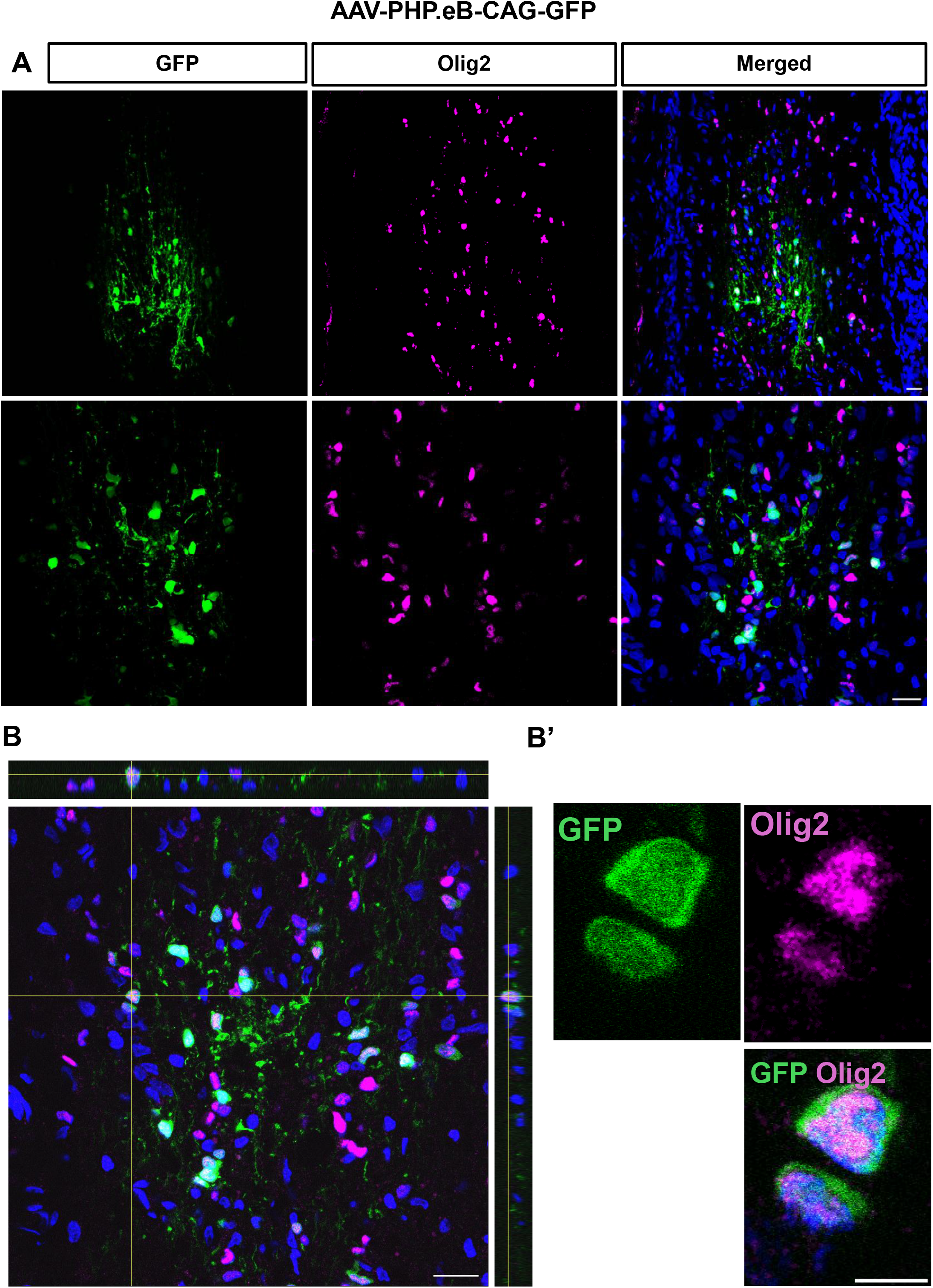
AAV-PHP.eB-CAG-GFP achieves strong gene delivery to Olig2 positive cells. (A) Representative confocal images from two optic nerves injected with AAV-PHP.eB-CAG-GFP showing GFP- and Olig2-positive cells. (B) Orthogonal projections demonstrate co-localization of GFP and Olig2 within individual cells. N= 4. Scale bars, 20 μm, and 5 μm in higher-magnification cropped image (B’).

**Figure 4.**
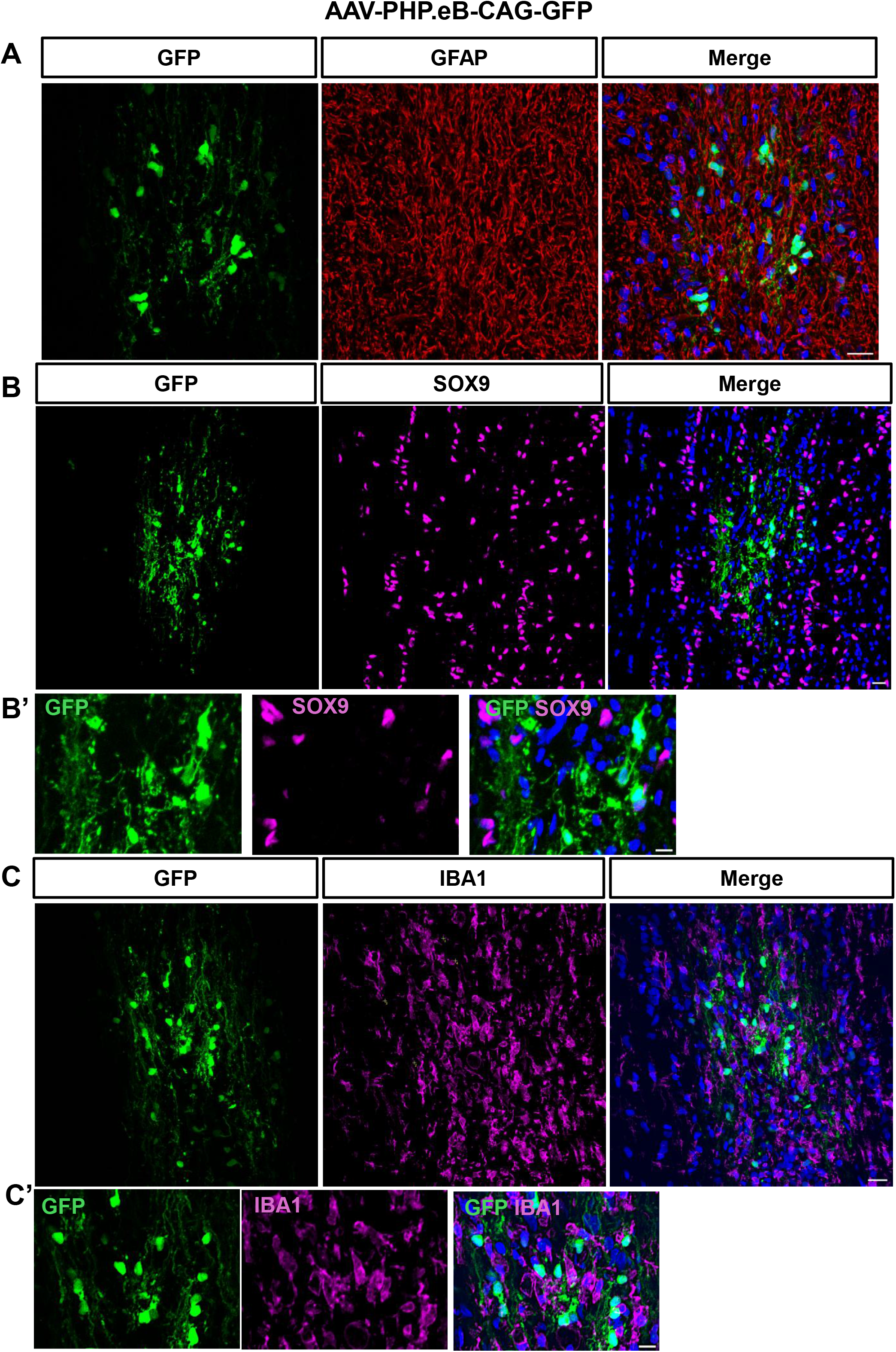
Assessment of AAV-PHP.eB-CAG-GFP–mediated gene delivery to astrocytes and microglia. Representative confocal images show GFP-positive (green) cells in optic nerve sections immunostained for GFAP (A), SOX9 (B), and IBA1 (C). (B’) A higher magnification image showing GFP and Sox9-immunoreactive cells. (C’) A higher magnification cropped image showing GFP and IBA1-immunoreactive cells. Nuclei are counterstained with DAPI (blue). N= 4. Scale bars, 20 μm, and 10 μm in the cropped images (B’ and C’).

**Figure 5.**
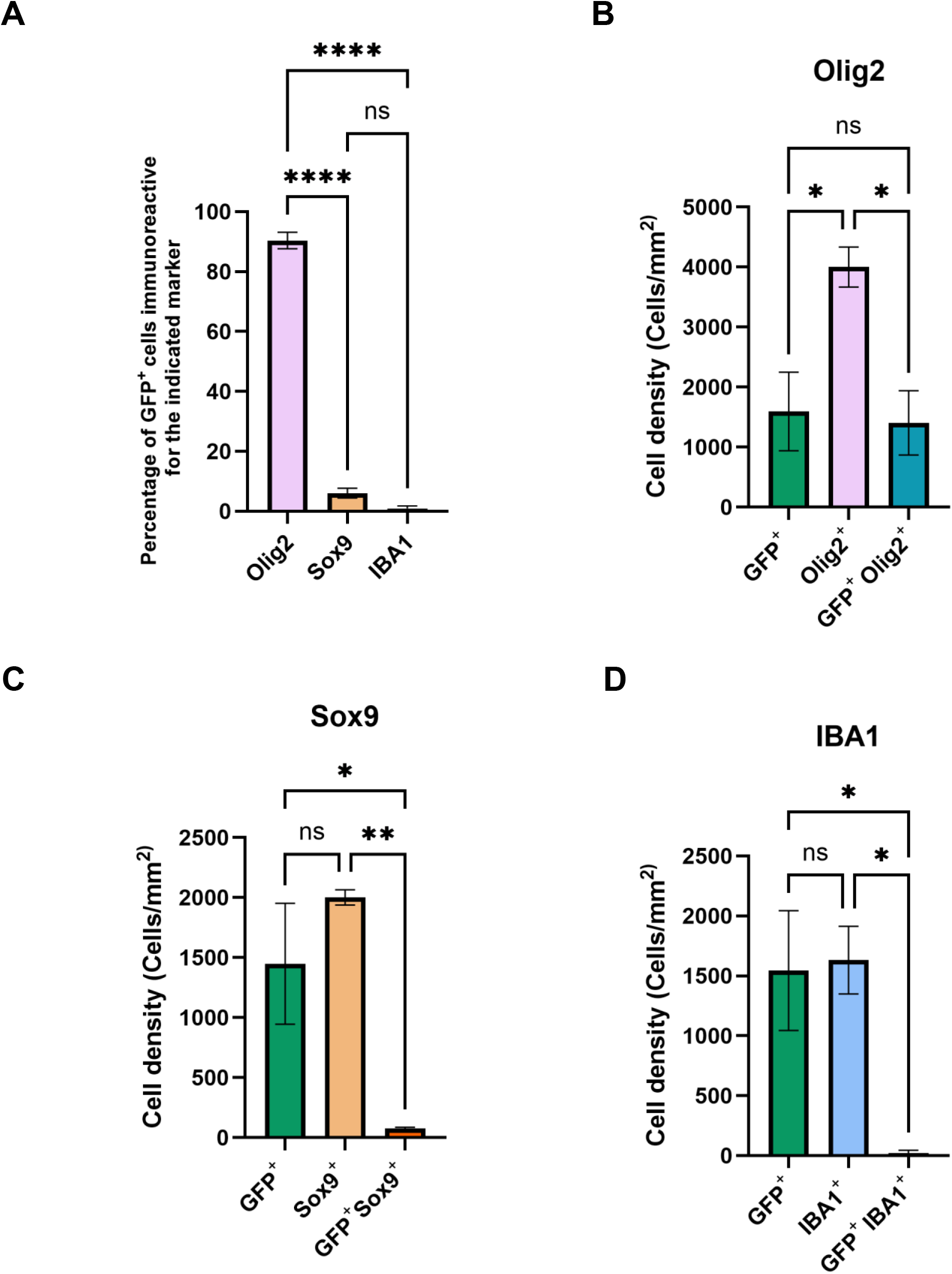
Quantification for cell-type-specificity and transduction efficiency of AAV-PHP.eB-CAG-GFP in the rat optic nerve. (A) Percentage of GFP^+^ cells colocalizing with each marker (Olig2, Sox9, and IBA1), indicating cell-type-specific transduction efficiency. (B-D) Quantification of the cell density (Cell/mm^2^) for Olig2, Sox9, and IBA1-positive cells, as well as GFP positive and double-positive cells (GFP^+^/marker^+^) in the rat optic nerve. One-way ANOVA followed by Tukey’s post hoc test revealed significantly higher transduction of Olig2^+^ cells compared with Sox9^+^ (****p < 0.0001) and IBA1^+^ cells (****p < 0.0001), while no significant difference was observed between Sox9^+^ and IBA1^+^ cells (ns). N= 3. Data are presented as mean ± SEM.

## DISCUSSION

This study, to our knowledge, is the first to systematically evaluate viral transduction of optic nerve cells using direct intra-nerve injections. Conventional ocular routes such as intravitreal and subretinal delivery have provided effective strategies for retinal gene transfer and even led to approved therapies for inherited retinal degeneration. A few studies have shown that intravitreal injection of adenovirus or AAV5 with a truncated GFAP promoter, gfaABC(1)D can deliver genes to glial cells in the optic nerve head.^31,32^ However, these approaches are largely ineffective at targeting optic nerve cells located farther from the eye.

Only a few studies have explored intra-optic nerve delivery of viral vectors ^33^. Binkley et al. demonstrated that AAV-eGFP injected into the rat optic nerve via a transconjunctival approach predominantly transduced cells in the optic nerve sheath, with limited gene expression in the parenchyma. However, the specific AAV serotype and promoter used in this study were not described.^33^

In the present study, we found that AAV2 and AAV5, even with ubiquitous promoters, achieved only modest transduction in the optic nerve. Lentivirus produced slightly higher transduction, but levels remained limited. This sparse transduction was somewhat unexpected, given the robust expression these viruses achieve at comparable titers in the brain or spinal cord.^34,35^ While the weak transduction could partly reflect the technical challenges of injecting into the smaller optic nerve, this is unlikely to be the sole explanation, as we observed strong transduction with AAV-PHP.eB.CAG under similar conditions.

Unlike the limited efficiency observed with AAV2 and AAV5, our results demonstrate that AAV-PHP.eB.CAG-GFP achieves reproducible transduction in the optic nerve, particularly in oligodendrocytes. Derived from AAV9, PHP.eB was engineered to enhance penetration through vascular and extracellular matrix barriers, enabling efficient gene delivery to both neurons and glia within white matter tracts. ^36^ Notably, while AAV-PHP.eB drove strong expression under the ubiquitous CAG promoter, transduction was weak when the GFAP promoter was used in rat optic nerve astrocytes. We speculate that the high efficiency of AAV-PHP.eB in targeting oligodendrocytes with the CAG promoter may reflect preferential receptor availability on these cells. Since local AAV-mediated transduction depends on serotype–receptor interactions (e.g., AAV2 binds heparan sulfate proteoglycans, AAV5 binds α2,3-linked sialic acid, and AAV9 binds N-linked galactose)^37–39^, it is plausible that AAV-PHP.eB exploit receptor profiles that are particularly abundant in oligodendrocytes, thereby enhancing its tropism in the optic nerve. Additionally, AAV-PHP.eB may efficiently enter optic nerve astrocytes; however, it is possible that the relatively weak activity of the GFAP promoter resulted in low transgene expression in this glial cell type. This may reflect the relatively weak and context-dependent nature of GFAP transcription, which is low in quiescent astrocytes and becomes strongly induced only under reactive or injury conditions. ^40–42^ Together, these findings indicate that conventional AAV2 and AAV5 may not be suited for direct optic nerve gene delivery in rats, whereas engineered variants like AAV-PHP.eB illustrates the potential of rationally evolved capsids to overcome intrinsic barriers in this tissue.

We note that intra-optic nerve injection would inherently cause a degree of local tissue damage due to direct penetration of the optic nerve by the injection pipette, even when using a fine tipped glass pipette and minimal injection volume. This mechanical disruption is unavoidable and is an important limitation of the technique. Despite this unavoidable local injury, we consistently observed viral transduction in glial cells surrounding the injection site. This indicates that the biological processes necessary for viral entry, gene expression, and cell type specific transduction remain intact in the injured microenvironment. Importantly, the goal of the present study was not to establish a damage free delivery method, but rather to determine whether effective and cell selective gene manipulation can be achieved within the adult optic nerve following direct injection. From this perspective, the presence of localized injury does not preclude the utility of the approach.

Instead, it provides a practical experimental context in which gene manipulation can still be achieved and leveraged to study distinct biological processes, such as glial responses, oligodendrocyte biology, and gene function in the injured or perturbed optic nerve. Indeed, controlled injury models are commonly used in CNS research, and targeted gene delivery within such contexts remains highly informative. Thus, intra-optic nerve injection can serve as a useful platform for mechanistic studies not necessarily as a procedural safety or therapeutic readiness. Additionally, future studies incorporating dedicated axonal integrity and functional assessments will be required to fully characterize injury extent and long-term safety.

The transduction of oligodendrocytes by AAV-PHP.eB-CAG in the adult optic nerve provides a valuable tool for investigating the role of these glial cells in CNS trauma and disease. Oligodendrocytes are critical for axonal myelination, metabolic support, and maintenance of neuronal integrity, and their dysfunction contributes to a variety of optic neuropathies. In conditions such as optic neuritis and multiple sclerosis, demyelination leads to impaired conduction and axon loss, while in glaucoma, oligodendrocyte stress may exacerbate retinal ganglion cell degeneration. Similarly, in optic nerve tumors such as those associated with neurofibromatosis type 1 (NF1), altered glial cell signaling can influence tumor microenvironments and neuronal survival. By enabling targeted gene manipulation in oligodendrocytes, AAV-PHP.eB-CAG allows researchers to interrogate molecular pathways that regulate oligodendrocyte survival, remyelination, and axon support. This vector also facilitates studies of axon regeneration following injury, as oligodendrocytes both provide inhibitory cues and metabolic support that influence regrowth. Collectively, these features make AAV-PHP.eB-CAG an ideal platform for mechanistic studies and preclinical intervention strategies aimed at preserving or restoring optic nerve function across diverse disease contexts. Its high specificity and efficiency in oligodendrocytes overcome limitations of previous viral vectors, opening new avenues for understanding glial contributions to CNS repair and neuroprotection.

We note that it is important to consider that direct needle injection into the optic nerve can itself induce tissue damage and local inflammation, potentially influencing experimental outcomes. Such injury could affect glial activation, axon integrity, or gene expression independently of the transgene being studied. Therefore, including appropriate control AAV injections using vectors expressing inert reporters like GFP, is critical when assessing gene-specific effects.

We note that, in this study, each treatment group included a minimum of two animals and up to four animals (n = 2–4 per group). We acknowledge that the sample size, particularly for groups with n of 2, is limited and represents a constraint of the current study. Thus, we acknowledge that the results should be viewed as exploratory and that future studies with larger cohorts may be required to validate and extend these observations.

In conclusion, our findings indicate direct optic nerve injection as a viable route for AAV-mediated gene transfer. However, we also note that the viral transduction remains largely localized rather than diffusely distributed throughout the entire nerve. By circumventing the limitations of intravitreal approaches, this method nonetheless provides access to axonal and glial compartments of the nerve and enables systematic evaluation of viral performance and future therapeutic impact. Although translational challenges remain, the ability to reproducibly target the optic nerve creates new opportunities for both mechanistic research and therapeutic development. Ultimately, direct nerve delivery complements existing ocular gene therapy strategies and may accelerate progress toward more effective research strategies interventions for optic neuropathies and other related CNS conditions.

