## Supplementary material for "Comparative Evaluation of Adeno-Associated Virus and Lentivirus Mediated Gene Transfer in Adult Rat Optic Nerve": Sup Figs.Legends

**Supplementary Figure 1. Schematic of optic nerve-targeted viral injection in the rat.** (A) Schematic illustration of the optic nerve injection setup using a nanoliter injector and a glass capillary to deliver viral vectors directly into the optic nerve. (B) Representative fluorescence image showing GFP expression (green) at the injection site in the optic nerve. Nuclei are counterstained with DAPI (blue). The injection site is indicated. Scale bars, 50 µm

**Supplementary Figure 2. Representative image of optic nerves with AAV5-CAG-GFP, AAV5-GFAP-GFP, or AAV-PHP.eB-GFAP injection**. GFP signal is shown on the green channel. A red channel without any immunostaining was included to identify autofluorescent cells, which exhibit overlapping signals in the red and green channels. Scale bars, 50 µm.
