## Supplementary figures and images for "Comparative Evaluation of Adeno-Associated Virus and Lentivirus Mediated Gene Transfer in Adult Rat Optic Nerve"

### Sup.Figs

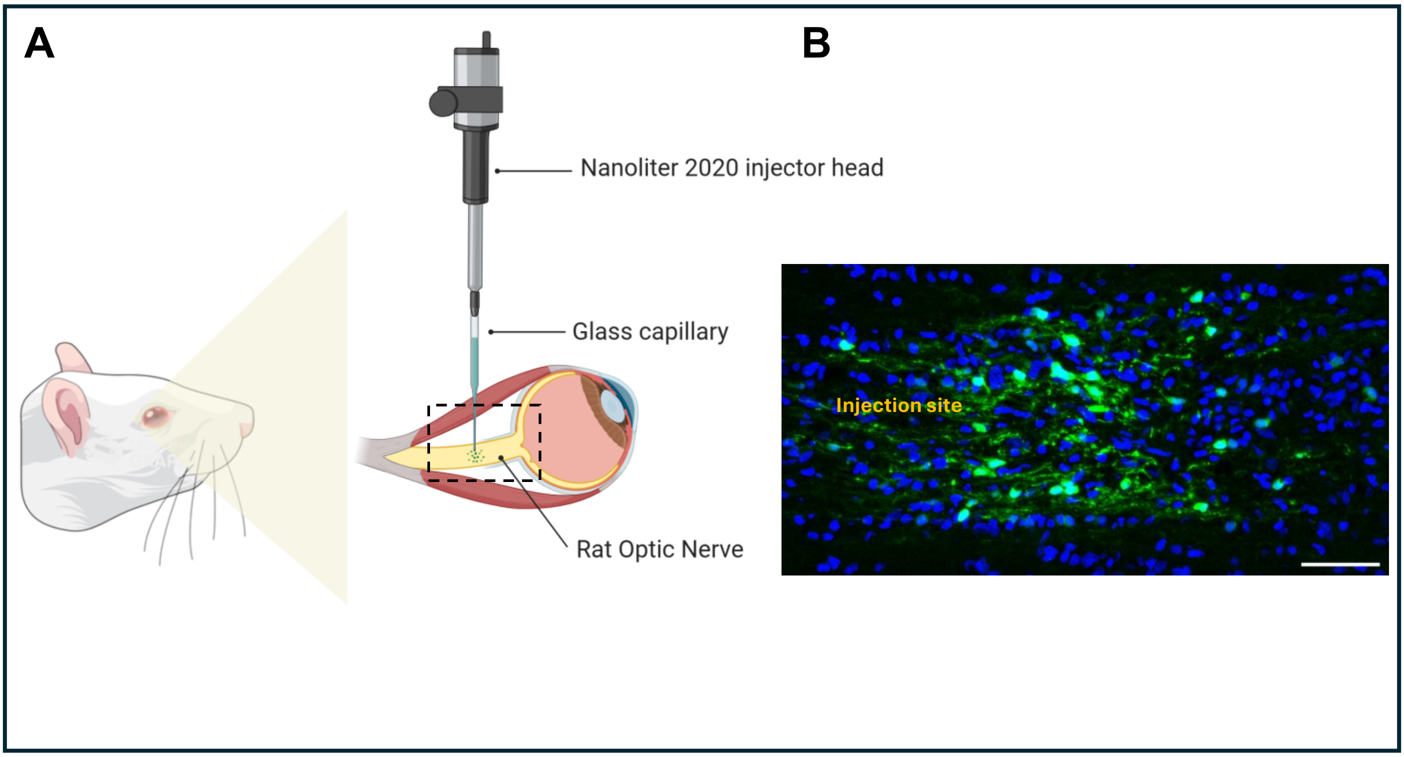


**Supplementary Figure 1**

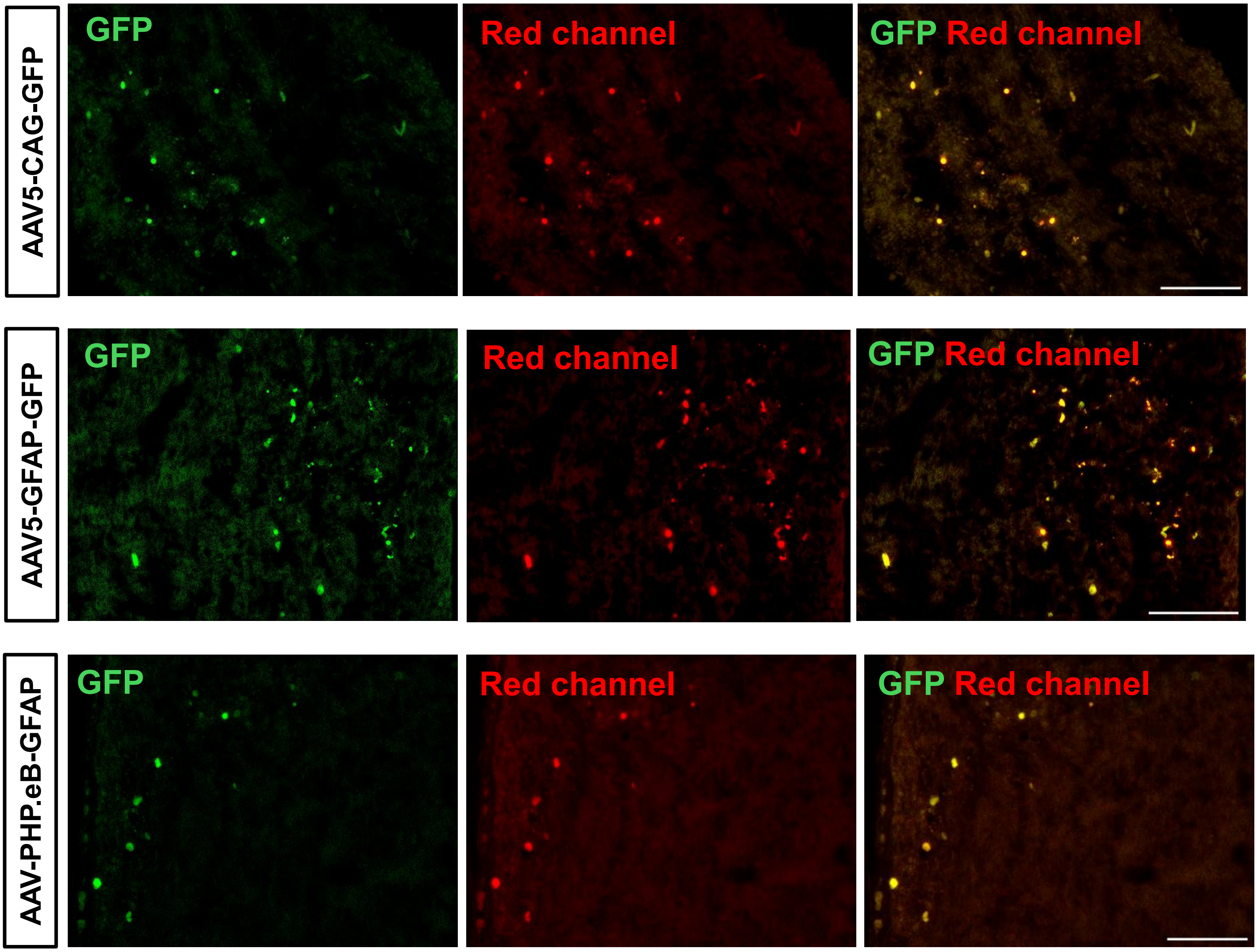


**Supplementary Figure 2**
